# The HIV-1 envelope cytoplasmic tail protects infected cells from ADCC by downregulating CD4

**DOI:** 10.1101/2025.06.06.658250

**Authors:** Alexandra Tauzin, Étienne Bélanger, Jérémie Prévost, Halima Medjahed, Catherine Bourassa, Frederic Bibollet-Ruche, Jonathan Richard, Beatrice H. Hahn, Andrés Finzi

## Abstract

HIV-1-mediated CD4 downregulation is a well-known mechanism that protects infected cells from antibody-dependent cellular cytotoxicity (ADCC). While CD4 downregulation by HIV-1 Nef and Vpu proteins has been extensively studied, the contribution of the HIV-1 envelope glycoprotein (Env) in this mechanism is less understood. While Env is known to retain CD4 in the endoplasmic reticulum (ER) through its CD4-binding site (CD4bs), little is known about the mechanisms underlying this process. Here we show that the cytoplasmic tail of Env is a major determinant in CD4 downregulation. This function is highly conserved as it was observed with nine different infectious molecular clones from four clades. The small but significant accumulation of CD4 at the surface of cells infected with Env cytoplasmic tail deleted viruses is sufficient to trigger Env to adopt a more “open” conformation. This prompted recognition of such HIV-1-infected cells by plasma from people living with HIV (PLWH) and several families of CD4-induced (CD4i) antibodies, leading to the elimination of these cells by ADCC. While cytoplasmic tail truncations are known to enhance Env expression at the cell surface, this did not fully explain the increased recognition of infected cells by CD4i antibodies and plasma from PLWH. Introduction of the CD4bs D368R mutation, which abrogates CD4 interaction, decreased Env recognition and ADCC. Overall, our results show that CD4 downregulation by the cytoplasmic tail of Env contributes to the protection of infected cells from ADCC.

**IMPORTANCE:** HIV-1-mediated CD4 downregulation is a central mechanism involved in the protection of infected cells from antibody-dependent cellular cytotoxicity. CD4 downregulation prevents the premature interaction between HIV-1 envelope glycoproteins (Env) and CD4, which would otherwise “open” Env and expose vulnerable epitopes recognized by CD4-induced antibodies present in the plasma from people living with HIV. While the mechanisms of CD4 downregulation by the viral accessory proteins Nef and Vpu have been elucidated, the function of Env in this process is less clear. Here we show that the cytoplasmic tail of Env plays an important role, thus contributing to the protection of infected cells from ADCC.

## OBSERVATION

CD4 downregulation is a central mechanism developed by HIV-1 to protect infected cells from antibody-dependent cellular cytotoxicity (ADCC) (1, 2). If CD4 is not downregulated, it interacts with HIV-1 envelope glycoproteins (Env) exposing otherwise occluded vulnerable epitopes recognized by CD4-induced (CD4i) antibodies present in plasmas from people living with HIV (PLWH) (3).

It is well established that HIV-1 uses several proteins to downregulate CD4 from the surface of infected primary CD4^+^ T cells. The accessory protein Nef targets CD4 molecules already present at the plasma membrane by engaging the clathrin-associated adaptor protein 2 (AP-2) complex, which accelerates CD4 endocytosis and degradation by the lysosomes (4–6). The accessory protein Vpu targets newly-synthesized CD4 in the endoplasmic reticulum (ER) by recruiting β-TRCP and targeting CD4 to the ER-associated degradation (ERAD) pathway (7, 8). Env has also been described to be involved in CD4 downregulation (9–11). While it has been shown that Env retains CD4 in the ER and that the CD4-binding site (CD4bs) of Env is required for this retention, little is known about additional determinants of Env that are involved in this process.

Env gets incorporated into virions through interaction of its cytoplasmic tail with the viral matrix (MA) protein (12), but other functions of this intracellular Env domain in viral replication are less understood (13). The Env cytoplasmic tail contains a membrane-proximal YxxΦ endocytosis motif responsible for Env internalization and recycling by the AP-2 clathrin dependent pathway (Figure 1A) (14). Mutations in this motif increase Env levels at the surface of infected cells, resulting in enhanced susceptibility to ADCC mediated by purified IgG from PLWH (15). However, it was previously shown that enhanced levels of Env at the surface of infected cells is not sufficient to render them more susceptible to ADCC by CD4i antibodies or plasma from PLWH (16). Instead, the transition of Env to a more “open” conformation, similar to the one induced by CD4 interaction, was also required (16). Therefore, we investigated whether the cytoplasmic tail of Env contributed to HIV-1-mediated CD4 downregulation to prevent this conformational change.

**Figure 1.**
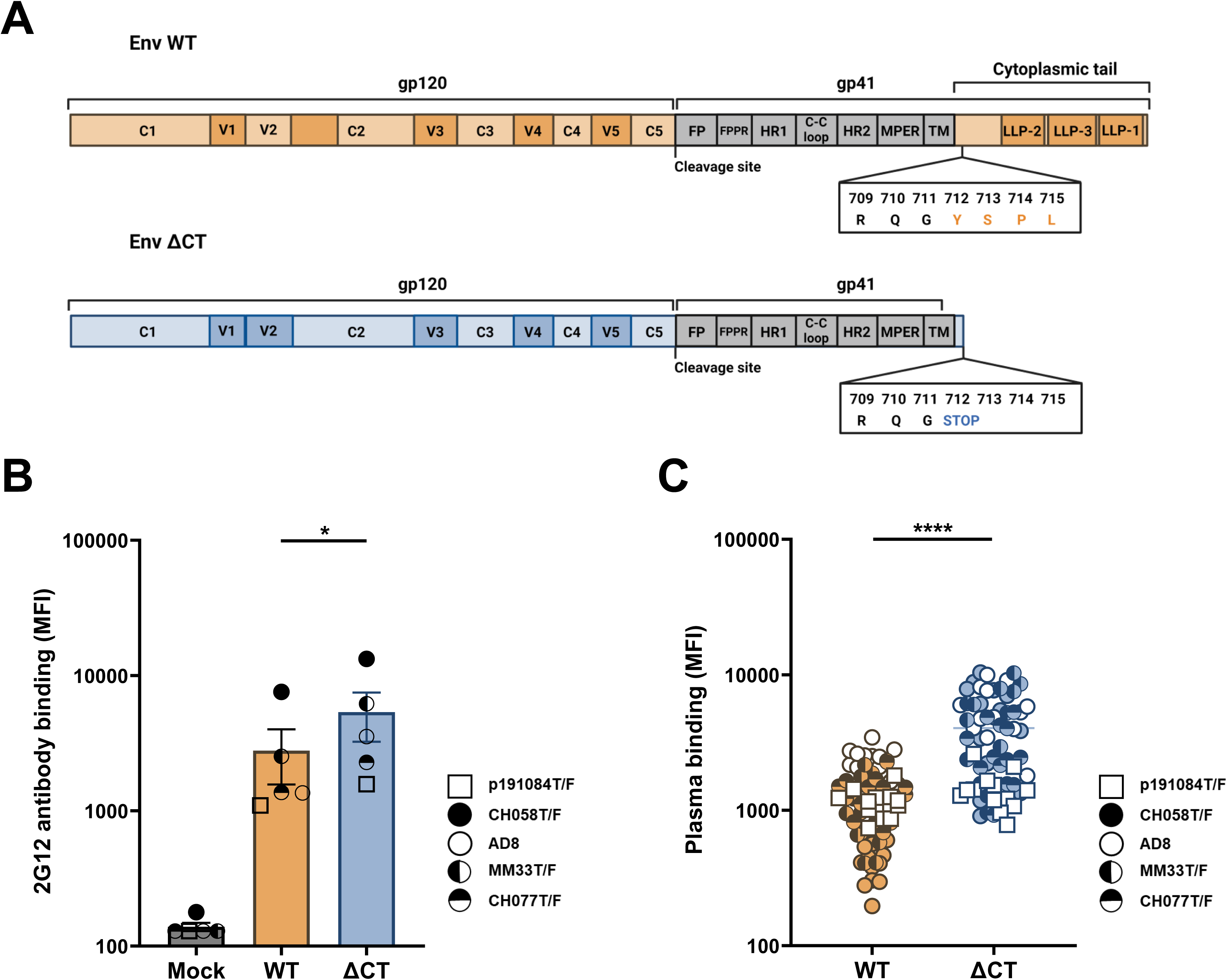
The HIV-1 Env cytoplasmic tail protects infected cells from detection by plasma from people living with HIV-1. (**A**) Schematic representation of the different domains of Env. C1-C5: constant domains; V1-V5: variable loops; FP: fusion peptide; FPPR: fusion peptide proximal region; HR: heptad repeat; MPER: membrane proximal external region; TM: transmembrane domain; LLP: lentivirus lytic peptides. (**B-C**) Primary CD4^+^ T cells were mock-infected or infected with viruses expressing either wild-type (WT) or cytoplasmic tail-truncated (ΔCT) Env from different clades. Two days post-infection, the cells were stained with (**B**) the conformation independent 2G12 monoclonal antibody (every symbol represents the mean of 2G12 binding for each IMC) or (**C**) 14 plasmas from PLWH. Shown are the median fluorescence intensities (MFI). Error bars indicate means ± SEM (∗p < 0.05 ∗∗∗∗; p < 0.0001). Statistical significance was tested using paired t tests, based on statistical normality.

We inserted a stop codon at position Y712 of the cytoplasmic tail, thereby truncating the tail just before the YxxΦ motif, in nine infectious molecular clones (IMCs) from four different clades (Figure 1A). IMCs expressing either wild type (WT) or cytoplasmic tail-deleted (ΔCT) Env were used to infect primary CD4^+^ T cells isolated from human peripheral blood mononuclear cells (PBMCs) from six HIV-negative donors. We first measured Env levels at the surface of infected cells using the conformation independent 2G12 antibody. As expected, we observed enhanced Env levels at the surface of Env ΔCT HIV-1-infected cells compared to their WT counterparts (Figures 1B and S1A). Deletion of the cytoplasmic tail also significantly improved Env recognition by plasma from PLWH (Table S1, Figures 1C and S1B), thus suggesting that Env is present in a more “open” conformation upon truncation of its cytoplasmic tail.

To evaluate whether this was the case, we probed these cells with four well-characterized CD4i antibodies recognizing the gp120 cluster A region (A32), the co-receptor binding site (17b), the V3 loop (19b) and the gp41 cluster I region (246D). In agreement with their CD4i nature, these antibodies poorly recognized cells infected with the WT virus but readily did so upon cytoplasmic tail truncation (Figure S2A). Since CD4 expression at the cell surface is one of the main factors affecting Env conformation, we next measured CD4 levels using the OKT4 antibody. Remarkably, deletion of the Env cytoplasmic tail led to a small, but significant, accumulation of CD4 at the surface of cells infected with the nine IMCs tested (Figures 2A-B and S3). These results indicated an important role of the Env cytoplasmic tail in CD4 downregulation.

**Figure 2.**
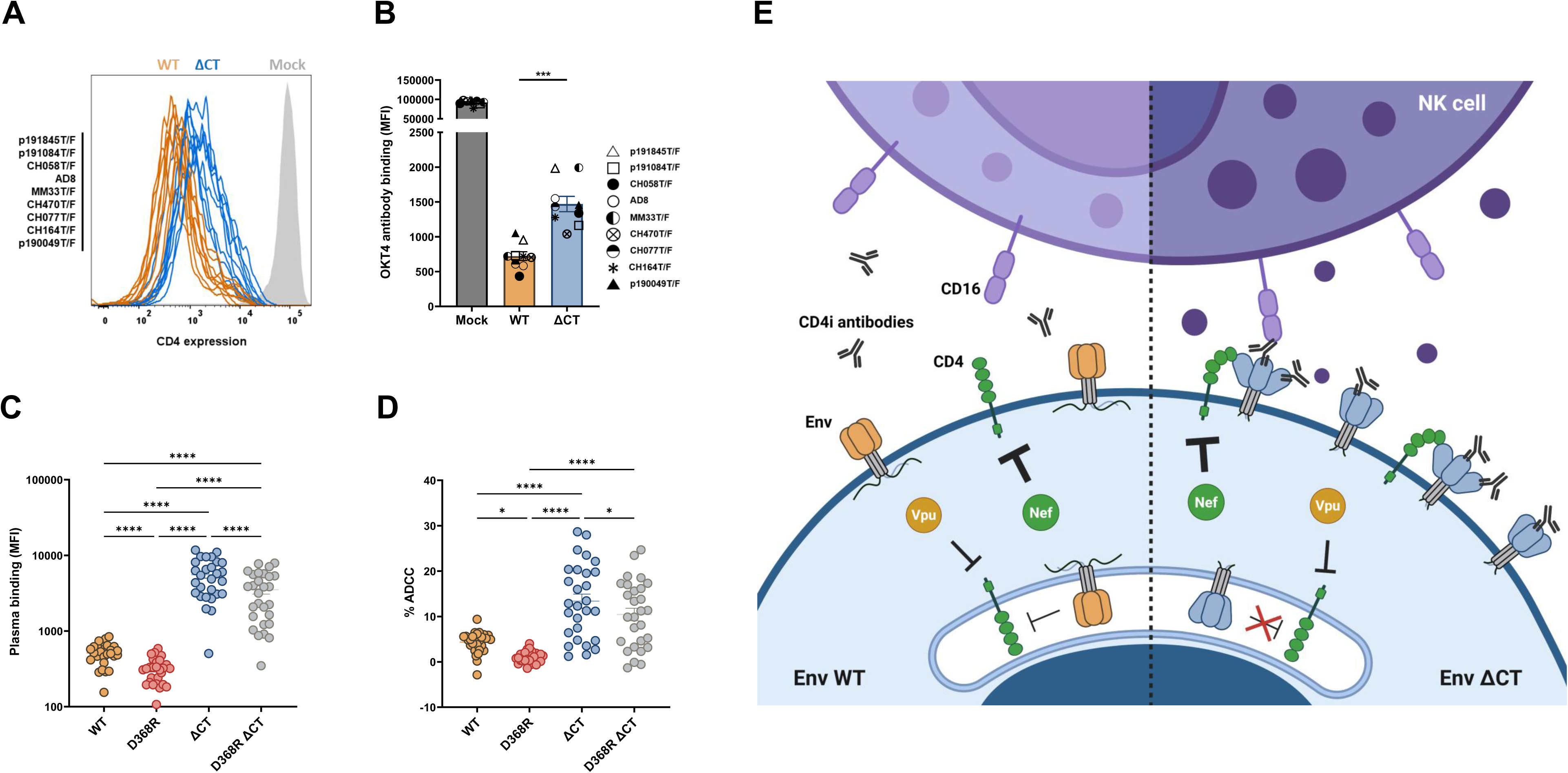
The HIV-1 Env cytoplasmic tail protects infected cells from ADCC by contributing to CD4 downregulation. (**A-B**) Primary CD4^+^ T cells were either mock-infected or infected with 9 different IMCs from 4 different clades, each expressing either WT or ΔCT Env. Two days post-infection, the cells were stained with the anti-CD4 OKT4 antibody to measure cell-surface CD4 levels. (**A**) Representative histograms showing surface CD4 expression on CD4^+^ T cells, either mock-infected (grey) or infected with indicated IMCs expressing WT (orange) or ΔCT (blue) Env. (**B**) Every symbol represents the mean of anti-CD4 OKT4 binding obtained for each IMC variant in at least 4 independent experiments. (**C-D**) Primary CD4^+^ T cells were infected with the transmitted/founder virus HIV-1_CH058T/F_ expressing either the WT, D368R, ΔCT or D368R ΔCT Env. Two days post-infection, (**C**) Env recognition and (**D**) ADCC-mediated elimination of infected cells by 28 plasmas from PLWH was measured by flow cytometry. Error bars indicate means ± SEM (∗p < 0.05; ∗∗∗p < 0.001; ∗∗∗∗p < 0.0001). Shown are the MFI and percentage of ADCC obtained. Statistical significances were tested using (**B**) paired t tests, (**C**) RM one-way ANOVA and (**D**) Friedman test, based on statistical normality. (**E**) Overview of the effect of cytoplasmic tail truncation on CD4 expression levels and Env conformation.

To evaluate whether this small CD4 accumulation at the surface of infected cells contributed to Env recognition by plasma from PLWH and CD4i antibodies, we introduced a mutation in the gp120 CD4-binding site (CD4bs) at position D368 of HIV-1_CH058T/F_ expressing the WT and ΔCT Env. The D368R mutation abrogates both Env-CD4 interaction, thereby keeping Env in its “closed” conformation (2), as well as the effect of the cytoplasmic tail on CD4 levels (Figure S4). Supporting a role for Env-CD4 interaction in the enhanced recognition of infected cells by CD4i antibodies and plasma from PLWH, insertion of this mutation decreased Env recognition by these ligands (Figures 2C and S2A). The effect of the D368R mutation was less pronounced for 17b and 19b antibodies (Figure S2A). These results are in agreement with previous studies showing that cytoplasmic tail deletions enable Env to sample downstream “open” conformations more readily, even in the absence of CD4 (17). However, A32 binding required Env-CD4 interaction as illustrated by the lack of recognition in both WT and ΔCT Env upon D368R introduction.

We next measured ADCC mediated by plasma from PLWH (Figure 2D) and CD4-induced antibodies (Figure S2B) against HIV-1_CH058T/F_ expressing WT or ΔCT Env in the presence and absence of the D368R mutation. As expected, little ADCC was observed with WT Env (Figure 2D). Cytoplasmic tail deletion increased ADCC mediated by plasma from PLWH and 17b, 246D and 19b; however, this was not the case for A32, suggesting that the tail truncation-induced conformational change was not sufficient to mediate efficient binding by this antibody (Figure 2D and S2B). Importantly, insertion of the D368R mutation decreased ADCC responses mediated by plasma from PLWH and CD4i antibodies.

In summary, our results support a model (Figure 2E) where deletion of the cytoplasmic tail leads to a better recognition and elimination of HIV-1-infected cells by plasma from PLWH due to (i) an increase in Env levels at the surface of infected cells (15) as well as (ii) an increase in CD4 levels which results in the exposure of otherwise occluded internal trimer epitopes. Thus, intrinsic “opening” of Env upon cytoplasmic tail truncation (17) contributes to this phenotype. Additional work is required to determine how the cytoplasmic tail of Env contributes to CD4 downregulation.

## ACKNOWLEDGMENTS

The authors thank the CRCHUM BSL3 and Flow Cytometry Platforms for technical assistance, Mario Legault from the FRQS AIDS and Infectious Diseases network and Madeleine Durand from CRCHUM for cohort coordination and clinical samples. We thank James Robinson (Tulane University Medical Center) for providing the plasmids to produce the A32 and 17b antibodies and Frank Kirchhoff (Ulm University Medical Center) for providing the infectious molecular clone (IMC) AD8 (Vpu+). The plasmids for the HC and LC of 2G12, 246D and 19b were obtained from the NIH AIDS reagents program. This study was supported by a CIHR Team Grant #197728, a project grant #451304 and a Canada Foundation for Innovation grant #41027 to A.F as well as support by the National institutes of Health to B.H.H. (R01 AI162646, UM1AI144371 and UM1AI164570). É.B. is a recipient of FRQS and CIHR master’s fellowships. J.P. was the recipient of a CIHR doctoral fellowship. A.T. was supported by a MITACS Elevation post-doctoral fellowship. Figures 1A and 2E were created with BioRender.com. The funders had no role in study design, data collection and analysis, decision to publish, or preparation of the manuscript.

## AUTHOR CONTRIBUTIONS

A.T., É.B., J.P., and A.F. conceived the study. A.T., É.B., J.P., J.R., B.H.H. and A.F. performed, analyzed, and interpreted the experiments. A.T. and É.B. performed statistical analysis. É.B., H.M., C.B., F.B-R., B.H.H., and A.F. contributed to unique reagents. H.M. and C.B. collected and provided clinical samples. A.T. and A.F. wrote the manuscript with input from others. All authors have read and agreed to the published version of the manuscript.

---

**Figure S1.**
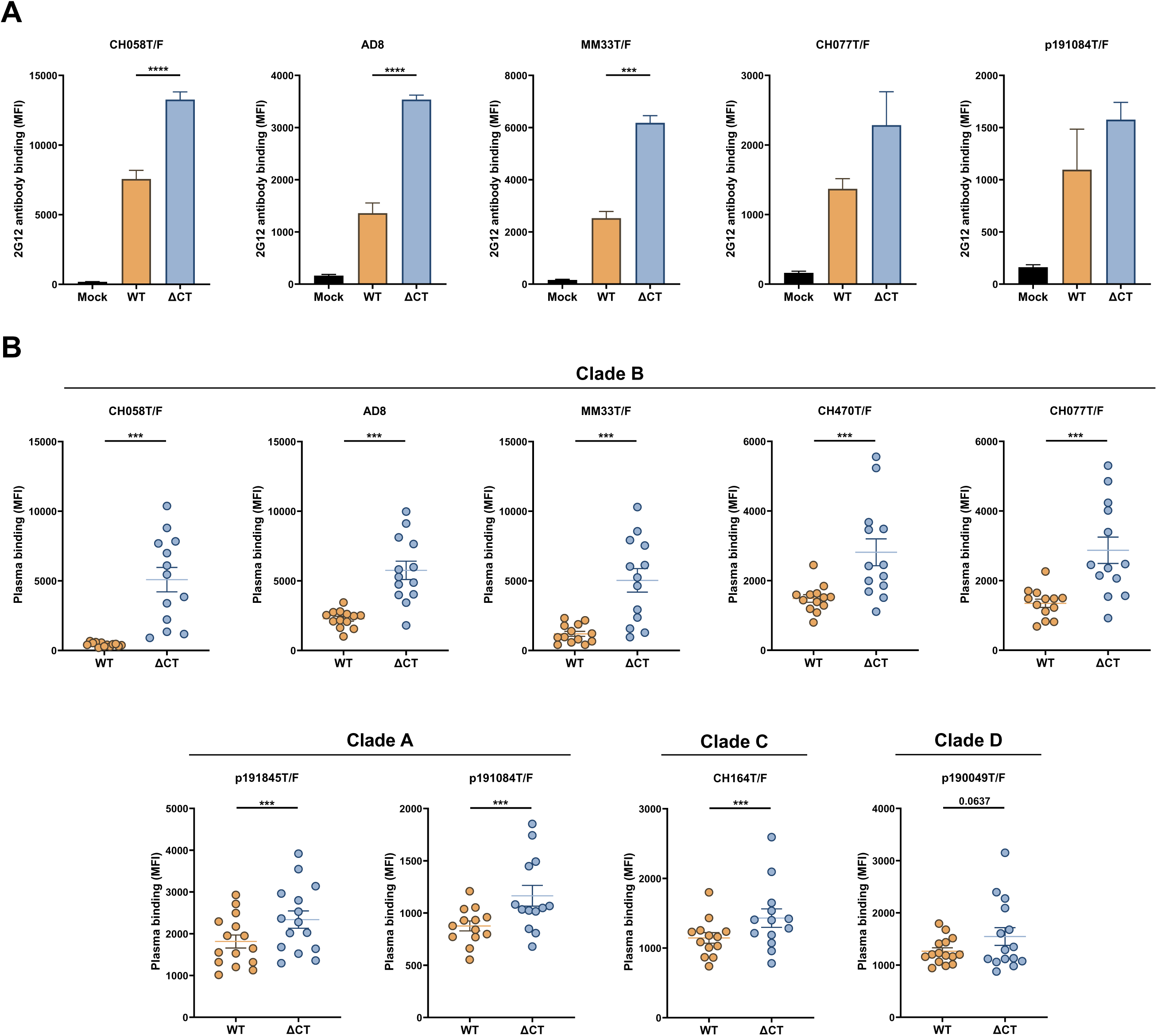
Env levels and recognition by plasma from PLWH at the surface of CD4^+^ T cells infected with WT or Env ΔCT viruses. Primary CD4^+^ T cells were mock-infected or infected with (**A**) 5 or (**B**) 9 primary viruses expressing either WT or ΔCT Env from (**A**) 2 or (**B**) 4 different clades. Two days post-infection, the cells were stained with (**A**) the conformation independant 2G12 antibody to measure Env levels at the surface of infected cells or (**B**) 14 plasma from PLWH. (**A**) The data shows the mean of at least 3 independent experiments. Error bars indicate means ± SEM (∗∗∗p < 0.001; ∗∗∗∗p < 0.0001). Statistical significance was tested using paired t tests or Wilcoxon matched-pairs signed rank test, based on statistical normality.

**Figure S2.**
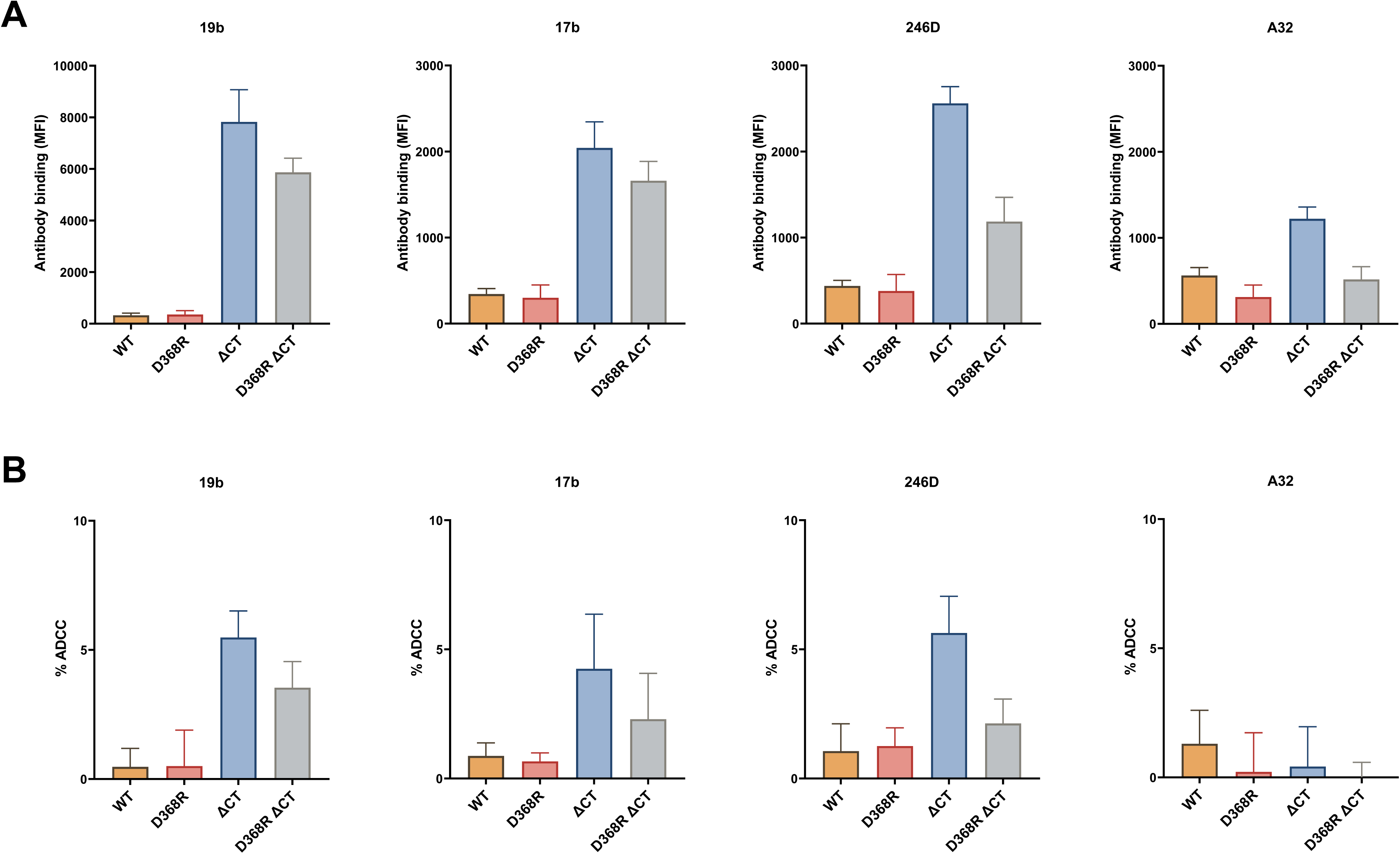
Deletion of Env cytoplasmic tail increases Env recognition and ADCC mediated by CD4-induced antibodies. (**A-B**) Primary CD4^+^ T cells were infected with the HIV-1_CH058T/F_ expressing either WT, D368R, ΔCT or D368R ΔCT Env. Two days post-infection, (**A**) Env recognition and (**B**) ADCC activity mediated by 19b, 17b, 246D or A32 antibodies were measured by cell surface staining and ADCC assay respectively.

**Figure S3.**
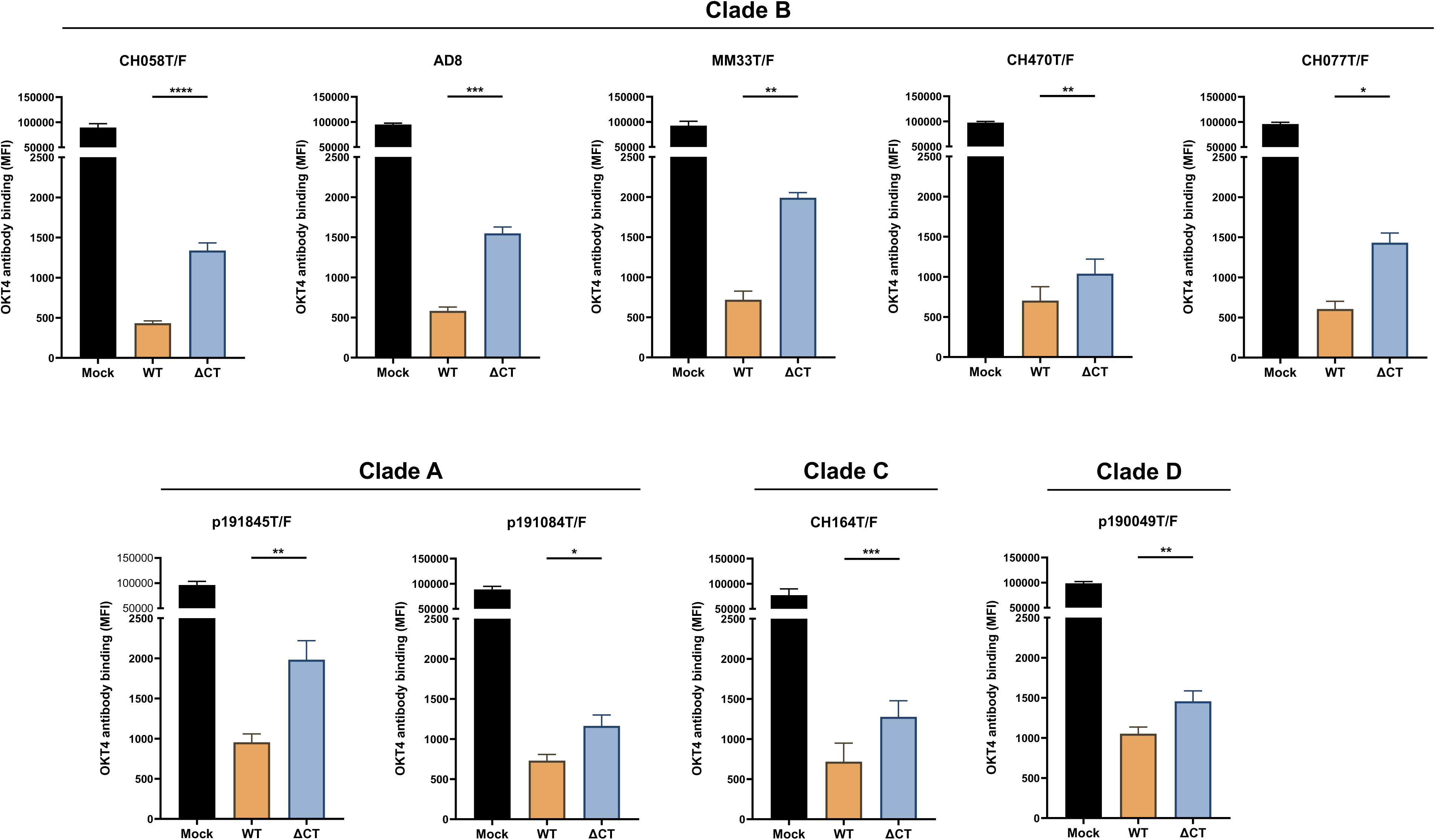
CD4 levels at the surface of CD4^+^ T cells infected with WT or Env ΔCT viruses. Primary CD4^+^ T cells were mock-infected or infected with 9 primary viruses expressing either WT or ΔCT Env from 4 different clades. Two days post-infection, the cells were stained with the anti-CD4 OKT4 antibody to measure CD4 levels at the surface of infected cells. The data shows the mean of at least 4 independent experiments. Error bars indicate means ± SEM (∗p < 0.05; ∗∗p < 0.01; ∗∗∗p < 0.001; ∗∗∗∗p < 0.0001). Statistical significance was tested using paired t tests or Wilcoxon matched-pairs signed rank test, based on statistical normality.

**Figure S4.**
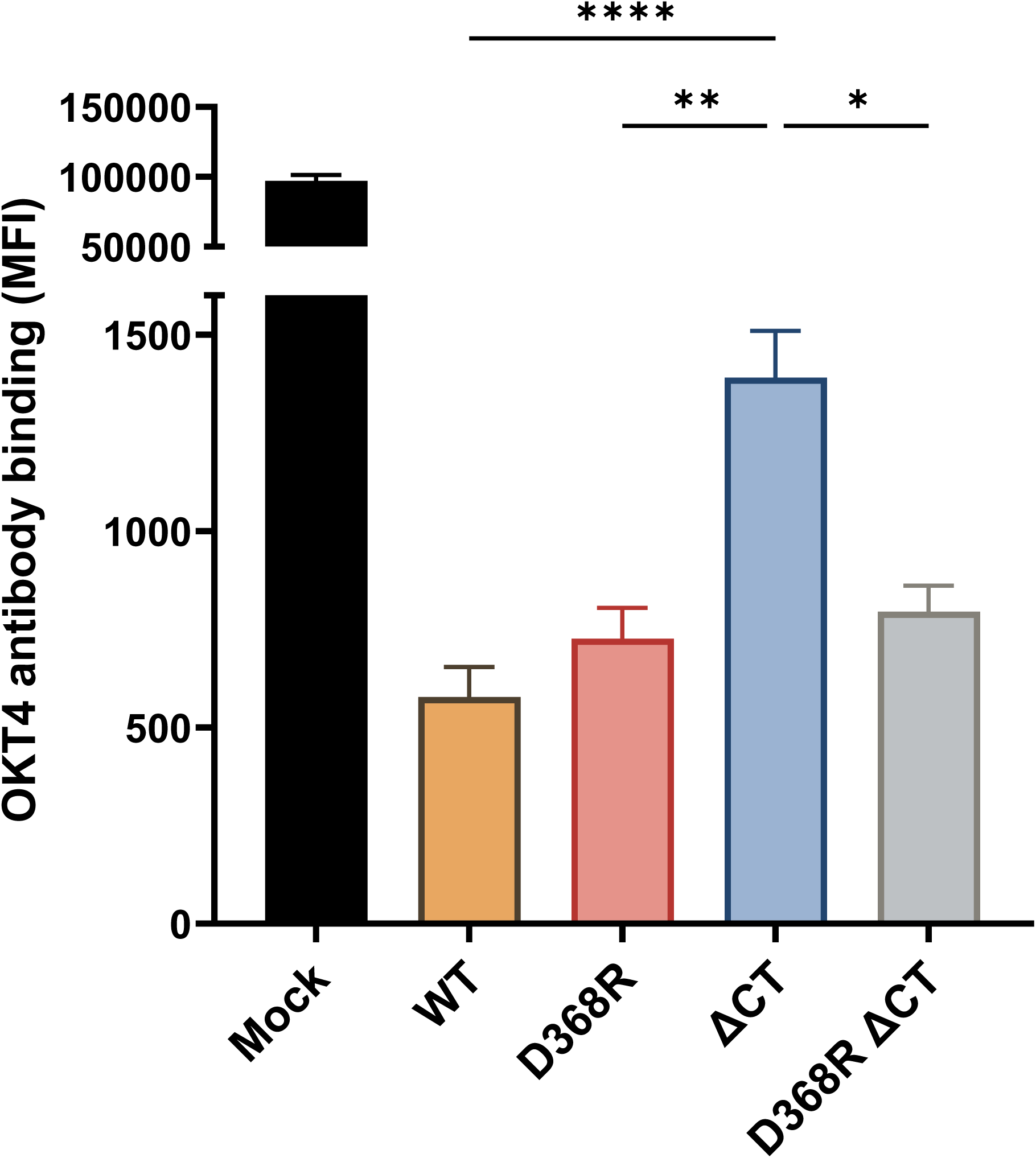
CD4 levels at the surface of CD4^+^ T cells infected with WT or Env ΔCT viruses harboring or not the D368R mutation. Primary CD4^+^ T cells were infected with the HIV-1_CH58T/F_ expressing either WT, D368R, ΔCT or D368R ΔCT Env. Two days post-infection, the cells were stained with the anti-CD4 OKT4 antibody to measure CD4 levels at the surface of infected cells. The data shows the mean of 10 independent experiments. Error bars indicate means ± SEM (∗p < 0.05; ∗∗p < 0.01; ∗∗∗∗p < 0.0001). Statistical significance was tested using Friedman test, based on statistical normality.

**Figure S5.**
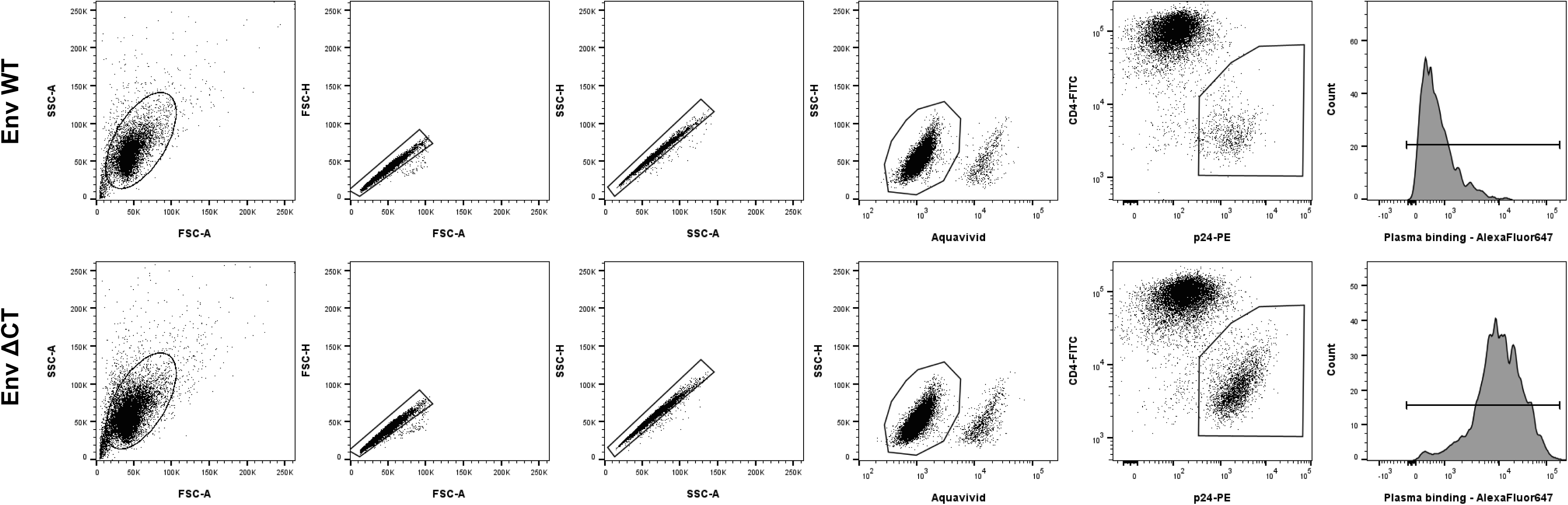
**Gating strategy for cell surface staining**. Representative flow cytometry gates to identify Env and CD4 levels at the surface of HIV-1 infected CD4^+^ T cells (top: WT; bottom: ΔCT Env). HIV-1 infected CD4^+^ T cells were stained with plasma or monoclonal antibodies and analyzed by flow cytometry. Cells were identified according to cell morphology by light-scatter parameters (first column) and excluding doublets cells (second and third columns). Cells were then gated on living cells (excluding the dead cells labeled with Aquavivid; fourth column). Finally, Env and CD4 binding by plasma or monoclonal antibodies was measured by the median of fluorescence of Alexa Fluor 647 (last column) in HIV-1 infected cells identified by gating on p24^+^CD4^low^ cells (fifth column).

**Figure S6.**
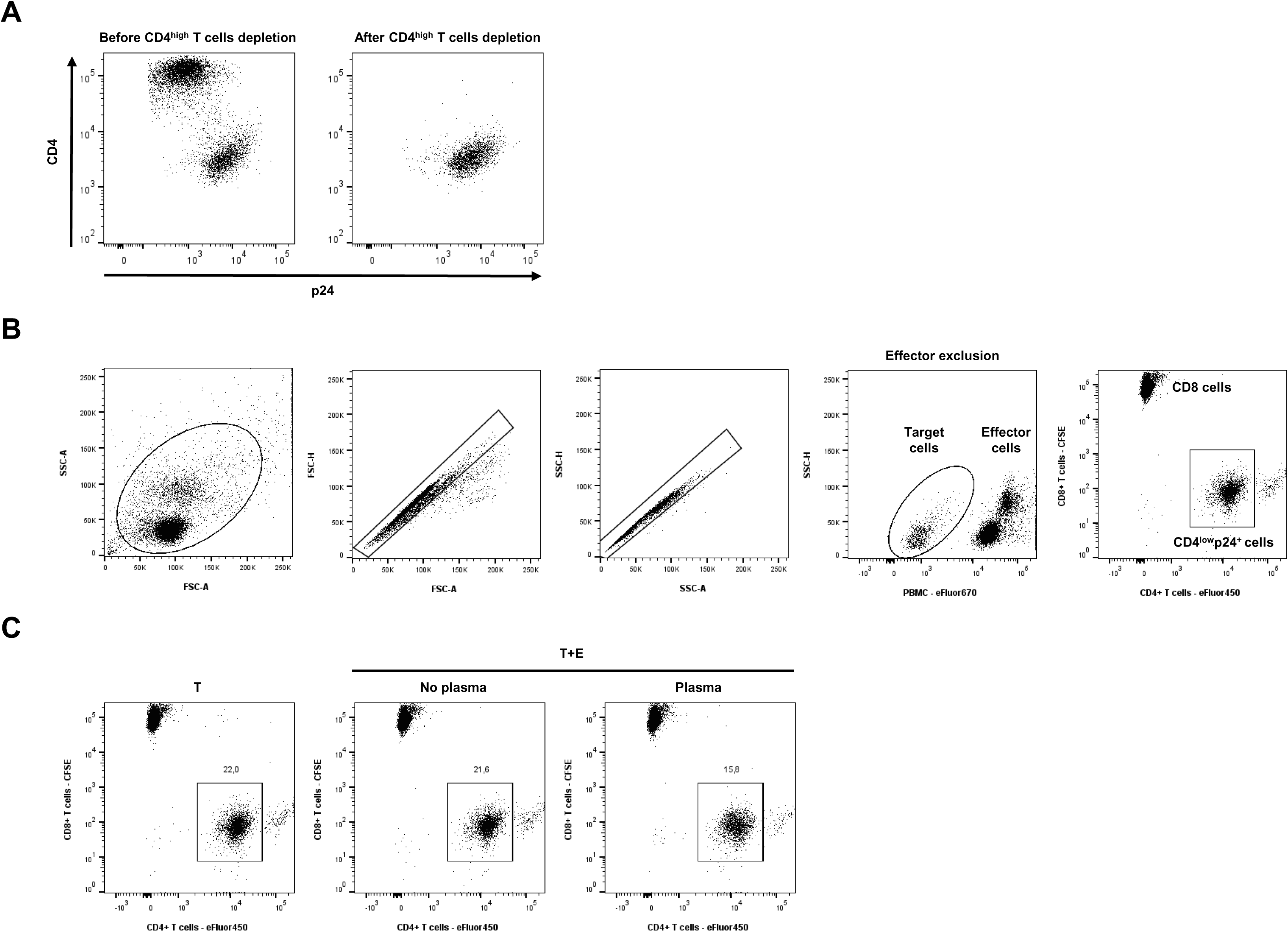
**Gating strategies for ADCC assay**. Primary CD4^+^ T cells were infected with HIV-1_CH58T/F_ expressing either WT, D368R, ΔCT or D368R ΔCT Env to perform ADCC assay. 48h post-infection, CD4^high^ uninfected bystander T cells coated with gp120 were depleted. (**A**) Representative flow cytometry gates to verify enrichement of productively-infected CD4^low^p24^+^ cells. The cells were stained with anti-CD4 OKT4 and anti-p24 monoclonal antibodies before and after removal of uninfected bystander CD4^high^ T cells. (**B**) Representative flow cytometry gates to measure ADCC activity. Total (target and effector cells) cells were identified according to cell morphology by light-scatter parameters (first column) and excluding doublets cells (second and third columns). Effectors cells were excluded by gating on eFluor670^-^ cells (fourth column). Finally, CD4^low^p24^+^ target cells were identified by gating on eFluor450^+^CFSE^-^ cells (last column). (**C**) Representative plots for the eFluor450^+^CFSE^-^ gate for Targets alone (T), and Targets with Effectors (T+E) in presence or absence of plasma from PLWH.

**Table S1:** Cohort of people living with HIV.

| Group | Sex |  | Age (years) | Years since ART initiation | Viral load (copies/mL) | CD4 count (cells/mm<sup>3</sup>) |
| --- | --- | --- | --- | --- | --- | --- |
|  | Female (n) | Male (n) |  |  |  |  |
| ART-treated | 3 | 18 | 51 (27-70) | 15.8 (0.1-23) | Undetectable | 699 (197-1250) |
| Untreated | 0 | 7 | 33 (29-51) | N/A | 6641 (132-372071) | 452 (263-570) |

\*Values displayed are medians, with ranges in parentheses.
ART: antiretroviral therapy.

## SUPPLEMENTAL MATERIALS AND METHODS

Supplemental material is available online.

Materials and methods have been previously reported in references (1–5) and are summarized below.

### Ethics statement

Written informed consent was obtained from all study participants and research adhered to the ethical guidelines of CRCHUM and was reviewed and approved by the CRCHUM institutional review board (ethics committee, approval number MP-02-2024-11734 and 11.062). Research adhered to the standards indicated by the Declaration of Helsinki. All participants were adult and provided informed written consent prior to enrollment in accordance with Institutional Review Board approval.

### Cell lines

HEK 293T human embryonic kidney cells (obtained from ATCC) were maintained in Dulbecco’s Modified Eagle Medium (DMEM) (Wisent) supplemented with 5% fetal bovine serum (FBS) (VWR) and 100 U/mL penicillin/streptomycin (Wisent) at 37°C under 5% CO_2._ FreeStyle 293F cells (Thermo Fisher Scientific) were grown in FreeStyle 293F medium (Thermo Fisher Scientific) to a density of 1 × 10^6^ cells/mL at 37°C under 8% CO_2_ with regular agitation (150 rpm).

### Primary cells

Human peripheral blood mononuclear cells (PBMCs) from six HIV-negative individuals (four males and two females, age range 31-67 years) were obtained by leukapheresis and Ficoll density gradient isolation and were cryopreserved in liquid nitrogen until further use. CD8^+^ T cells, used as control target cells in the ADCC assay, were isolated by negative selection following manufacturer’s instruction (StemCell Technologies, Cat #17953) and cultured at 2 million of cells/mL overnight in RPMI 1640 (Thermo Fisher Scientific) complete medium supplemented with 20% FBS and 100 U/mL penicillin/streptomycin. CD4^+^ T cells were purified from resting PBMCs by negative selection using immunomagnetic beads per the manufacturer’s instructions (StemCell Technologies, Cat #17952), were activated with phytohemagglutinin-L (PHA-L, 10 μg/mL) for 48h and then maintained in RPMI 1640 complete medium supplemented with 20% FBS, 100 U/mL penicillin/streptomycin and recombinant IL-2 (rIL-2, 100 U/mL). The cells were maintained at 37°C under 5% CO_2._

### Antibodiy production and purification

FreeStyle 293F cells were transfected with plasmids expressing the light (LC) and heavy (HC) chains of anti-Env monoclonal antibodies using ExpiFectamine 293 transfection reagent, as directed by the manufacturer (Thermo Fisher Scientific). One week later, the cells were pelleted and discarded. The supernatants were filtered (0.22-μm-pore-size filter), and the antibodies were purified by protein A affinity columns, as directed by the manufacturer (Cytiva). The recombinant protein preparations were dialyzed against phosphate-buffered saline (PBS) and stored in aliquots at −80°C. To assess purity, recombinant proteins were run on SDS-PAGE in the presence or absence of β-mercaptoethanol and stained with Coomassie blue.

### Antibodies and plasmas

Plasma from PLWH, obtained from the FRQS-AIDS and Infectious Diseases Network (the Montreal Primary HIV Infection Cohort, Table S1)(6, 7), were collected, heat-inactivated for 1 h at 56°C and stored at −80°C until ready to use in subsequent experiments. Plasma were used at a 1:1000 dilution in cell surface staining and ADCC assays.

The following monoclonal antibodies were used at 5 ug/mL to detect Env expression at the cell surface of HIV-1-infected cells: anti-gp120 outer domain 2G12 antibody (plasmids for HC and LC obtained from the NIH AIDS Reagent Program); anti-cluster A A32 antibody (plasmids for HC and LC kindly provided by James Robinson); anti-CoRBS 17b antibody (plasmids for HC and LC kindly provided by James Robinson); anti-gp41 cluster I 246D antibody (plasmids for HC and LC obtained from the NIH AIDS Reagent Program); anti-V3 loop 19b antibody (plasmids for HC and LC obtained from the NIH AIDS Reagent Program). The anti-CD4 monoclonal antibody (Clone OKT4, eBioscience, Catalog #14-0048-82) was used to detect CD4 expression in cell surface staining (1 ug/mL). Alexa Fluor-647-conjugated goat anti-human IgG (Thermo Fisher Scientific, Cat #A-21445) was used as secondary antibody to detect anti-Env antibodies and plasma binding by flow cytometry (2 µg/mL). Alexa Fluor-647-conjugated goat anti-mouse IgG (Thermo Fisher Scientific, Cat # A-21235) was used as secondary antibody to detect mouse anti-human CD4 monoclonal antibodies binding by flow cytometry (2 ug/mL). The FITC anti-human CD4 antibody (Clone OKT4, Biolegend, Cat # 6604667) (1:500 dilution) and the PE-conjugated anti-HIV p24 antibody (Clone KC57-RD1, Beckman Coulter, Cat # 6604667) (1:100 dilution) were used to identify productively-infected cells as previously described (8) in Env cell-surface staining.

### Plasmids and proviral constructs

The vesicular stomatitis virus G (VSV-G)-encoding plasmid was previously described (7). The infectious molecular clones (IMCs) of HIV-1_AD8_ as well as the transmitted/founder viruses HIV-1_CH058T/F_, HIV-1_CH470T/F_, HIV-1_CH077T/F_, HIV-1_CH164T/F_, HIV-1_MM33T/F_, HIV-1_p191845T/F_, HIV-1_p191084T/F,_ HIV-1_p190049T/F_ were previously described (10–15). The IMCs expressing ΔCT Env were generated by inserting a STOP codon in the YxxΦ internalization motif (at position Y712, according to HxBc2 numbering) by site-directed mutagenesis using the QuikChange II XL site-directed mutagenesis protocol (Agilent, Cat # 200521). The IMC HIV-1_CH058T/F_ expressing D368R and D368R ΔCT Env were generated by inserting the D368R mutation into HIV-1_CH058T/F_ expressing WT or ΔCT Env. The presence of the desired mutations was verified by automated DNA sequencing.

### Viral production *and in vitro* infections

VSV-G-pseudotyped HIV-1 viruses were produced by co-transfection of HEK 293T cells with the HIV-1 IMC proviral constructs and the VSV-G-encoding vector at a ratio of 3:2 using the polyethylenimine (PEI) method. Two days post-transfection, cell supernatants were harvested, clarified by low-speed centrifugation (300 × g for 5 min), and concentrated by ultracentrifugation at 4°C (100,605 × g for 1h) over a 20% sucrose cushion. Pellets were resuspended in fresh RPMI 1640 complete medium, aliquoted and stored at −80°C until use. VSV-G-pseudotyped HIV-1 viruses were then used for *in vitro* infection. Activated primary CD4^+^ T cells were spinoculated with the virus at 800 × *g* for 1h in 96-well plates at 25°C and then incubated 48h at 37°C. All viral productions were titrated on primary CD4^+^ T cells to achieve similar levels of infection (around 12-20% of infected cells).

### Flow cytometry analysis of cell-surface staining

Forty-eight hours post-infection, mock-infected and HIV-1-infected primary CD4^+^ T cells were collected, washed with PBS and transferred in 96-well V-bottom plates. The cells were then incubated for 45 min at 37°C with plasma (1:1000 dilution) or the 2G12, 19b, 17b, 246D or A32 monoclonal antibodies (5 µg/mL) or anti-CD4 OKT4 antibody (1 ug/mL).

Cells were then washed twice with PBS and stained with appropriated anti-IgG Alexa Fluor 647-conjugated secondary antibody (2 μg/mL), FITC-conjugated mouse anti-human CD4 Antibody (Biolegend, 1:500 dilution) and LIVE/DEAD viability dye (Thermo Fisher Scientific, Cat #L34957, 1:1000 dilution) for 20 minutes at room temperature. Cells were then washed twice with PBS and fixed in a 2% PBS-formaldehyde solution. The cells were then permeabilized using the Cytofix/Cytoperm Fixation/Permeabilization Kit (BD Biosciences, Cat #554714) and stained intracellularly using PE-conjugated mouse anti-p24 monoclonal antibody (clone KC57, 1:100 dilution). Samples were acquired on a Fortessa cytometer (BD Biosciences), and data analysis was performed using FlowJo v10.5.3. Env and CD4 levels at the surface of infected cells were measured by the median of fluorescence of Alexa Fluor 647 in productively infected cells determined by gating on living p24^+^ CD4^low^ cell population as previously reported (8) (Figure S5).

### ADCC assay

ADCC activity was measured 48h post-infection using a modified FACS-based infected cell elimination assay. It has been well established that soluble gp120 sheds from productively-infected cells and coats uninfected bystander cells; this substantially affects ADCC readings (4).To minimize this confounding factor and a potential effect of cytoplasmic tail truncation on gp120 shedding, the CD4^high^ T cells, which represent the uninfected bystander cells coated with gp120, were depleted from the target cell population using the Dynabeads® CD4^+^ positive selection kit (Thermo Fisher Scientific, Cat #11145D) at a ratio of 25 μl of beads per million cells as previously reported (3, 5). This procedure enriched the productively-infected CD4^low^p24^+^, as shown in Figure S6A. To specifically assess ADCC-mediated killing of enriched productively-infected CD4^low^p24⁺ cells, autologous resting CD8⁺ T cells, which are not susceptible to gp120 coating and ADCC, were included as control target cells (Figure S6B). The productively-infected CD4^low^p24^+^ cells were stained with the cell proliferation dye eFluor450 (Thermo Fisher Scientific, cat # 65-0842-90) while resting autologous CD8^+^ T cells were stained with the cell proliferation dye CFSE (Thermo Fisher Scientific, cat # C34554). The CD4^low^p24^+^ cells were then mixed with CD8^+^ T cells at a 1:3 ratio. Resting autologous PBMCs stained with the cell proliferation dye eFluor670 (Thermo Fisher Scientific, cat # 65-0840-90) were used as effectors cells. The target cell mix (CD4^low^p24^+^ cells and CD8^+^ T cells) was then co-cultured with autologous PBMCs (Effector: Target ratio of 10:1) in 96-well V-bottom plates in the presence of plasma from PLWH (dilution 1:1000) or the 19b, 17b, 246D or A32 monoclonal antibodies (5μg/mL) for 5h at 37°C. After incubation, cells were washed once with PBS and fixed in a 2% PBS-formaldehyde solution. Samples were acquired on a Fortessa cytometer (BD Biosciences), and data analysis was performed using FlowJo v10.5.3. The percentage of ADCC was calculated as follows: [(% of eFluor450^+^ cells in Targets plus Effectors) − (% of eFluor450^+^ cells in Targets plus Effectors plus plasma)/(% of eFluor450^+^ cells in Targets) × 100] (Figure 6C).

## QUANTIFICATION AND STATISTICAL ANALYSIS

Statistics were analyzed using GraphPad Prism version 10.2.0. Every data set was tested for statistical normality and this information was used to apply the appropriate (parametric or nonparametric) statistical test. Statistical details of experiments are indicated in the figure legends. p values < 0.05 were considered significant; significance values are indicated as *p < 0.05, **p < 0.01, ***p < 0.001, ****p < 0.0001.

